# Mapping Global Causal Responses to Noninvasive Modulation of Genetically and Spatially Targeted Neural Populations with Sonogenetic-fPET

**DOI:** 10.1101/2024.12.10.627860

**Authors:** Yaoheng Yang, Charlotte Weixel, Zhongtao Hu, Yimei Yue, Isaac Wang, Lu Xu, Hong Chen

## Abstract

Despite significant progress in brain circuit mapping over recent decades, a major challenge remains: no method currently allows for the noninvasive modulation of genetically and spatially defined neural populations while simultaneously monitoring their global effects throughout the brain and body. Here, we present sonogenetic-fPET, a technique that integrates sonogenetics with [^18^F]-2-fluoro-2-deoxy-D-glucose functional positron emission tomography (FDG-fPET) to overcome this challenge. Sonogenetics enables noninvasive, spatially targeted modulation of neurons genetically engineered to express the ultrasound-sensitive ion channel TRPV1, while FDG-fPET captures glucose metabolic changes triggered by this stimulation across the brain and body. We demonstrate the effectiveness of this technique by targeting neurons in the dorsal striatum, showcasing its capability to map global network responses to specific neuronal activation. Incorporating an acoustic hologram for sonogenetics further enables flexible modulation of different brain regions within a single mouse while concurrently mapping the resulting network activity. In summary, sonogenetic-fPET offers a tool for dissecting the global responses of the brain and body to the noninvasive modulation of genetically and spatially defined neuronal populations.

## Main

Understanding functional neural networks is crucial for advancing neuroscience and developing new treatments for brain diseases. While significant progress has been made in dissecting brain circuits, our progress is limited in connecting cell-type-specific neuromodulation with large-scale mapping of functional neural networks. This connection is critical for revealing the causality of neural functions and enabling the development of circuit-specific precise treatments for brain diseases. Opto-fMRI, which integrates optogenetics and functional MRI, has provided a method for activating genetically defined neurons at the site of optical fiber insertion and capturing the causal effects on neural circuits throughout the brain^1–3^. However, Opto-fMRI is constrained by the invasive nature of optogenetics and the limited capability to map global causal responses in the brain and body^1–3^. Recent advancements in sonogenetics and functional PET (fPET) laid the groundwork for the development of sonogenetics-fPET, a novel technique that enables the noninvasive modulation of genetically and spatially defined neuronal populations while simultaneously mapping global network response in both the brain and body.

Genetic-based neuromodulation approaches, such as the widely used optogenetics^4^ and chemogenetics^5^, provide cell-type specific neuromodulation. However, optogenetics typically requires surgical procedures to insert implants in the brain to deliver light to targeted brain regions^6^, particularly those deep in the brain, while chemogenetics lacks temporal resolution^5^. Although optogenetics stimulation offers rapid temporal control of neural activity, the need for implants restricts its application to typically one target site per animal^6,7^. These implants also pose risks of damaging neurons and non-neuronal cells, compromising the integrity of the neural circuit^8^. Significant efforts have been made to develop deep-focusing optical technologies without implants, such as multi-photon optogenetics^9,10^, adaptive optics^11^, wavefront shaping^12^, and acoustic-optic lenses^13^. However, achieving light focusing on deep tissue remains technically challenging due to the fundamental physical limitations of electromagnetic waves^14^. Exciting developments in implant-free deep brain optogenetics using red-shifted opsins activable by transcranial red light illumination have emerged^8^. Red light can penetrate deep into the mouse brain, but it loses the capability to target a specific brain region due to light scattering. Sonogenetics, which combines ultrasound with genetic engineering, offers unique capabilities for noninvasive, spatially targeted, and cell-type-specific neuromodulation^15–18^. Ultrasound can penetrate the intact skull to target deep brain regions and can be precisely focused on a small area, ranging from submillimeter to millimeter scale^19,20^, comparable to the focal size of an implanted optical fiber. Sonogenetics has made significant progress, evolving from its use in *Caenorhabditis elegans* to mice. Ion channels, such as MscL^21–23^, Prestin^24,25^, hsTRPA1^26^, and TRPV1^16–18^ have been utilized as sonogenetics actuators, with early studies demonstrating successful modulation of motor circuits in anesthetized mice. The development of wearable ultrasound devices enabled sonogenetic neuromodulation in freely moving mice^16^. We were the first to achieve modulation of a deep brain site, the striatum, in freely moving mice using TRPV1-sonogenetics^16^. Recently, we integrated acoustic hologram into sonogenetics, enabling flexible ultrasound beamforming to target any single location or multiple locations within the mouse brain^18^. These advancements in sonogenetics offer the unique capability to achieve noninvasive, flexible, and cell-type-specific neuromodulation in the whole brain.

Technological advancements have also accelerated our ability to map neural circuits, evolving from invasive single-point recording methods like optical-photometry and electrophysiology to noninvasive whole-brain imaging technologies like fMRI^27,28^ and fPET^29–32^. fMRI relies on the analysis of blood-oxygenated-level-dependent (BOLD) signal fluctuations due to neurovascular coupling ^27^. It can identify functional connectivity by detecting concurrent activations in different brain regions during sensory stimuli or cognitive tasks. fPET uses radiotracers, particularly [^18^F]-2-fluoro-2-deoxyglucose (FDG), to measure regional brain glucose metabolism, to reflect brain activity^29–32^. fPET has been instrumental in mapping brain activity and understanding the connectivity of neural circuits in response to various stimuli^33^. Recently, with further advancements and a deeper mechanical understanding, fPET has demonstrated its unique capability to image whole-body metabolic activity^34^, surpassing fMRI, which is primarily confined to the brain. These advancements in fPET provide a unique opportunity to simultaneously map neural circuits and delineate their functional outputs in peripheral tissues and organs.

Building on the advancements in sonogenetics and fPET, we developed sonogenetic-fPET to map the global responses of the brain and body to the modulation of genetically and spatially defined neuronal populations in a noninvasive manner. Using the basal ganglia circuit as an example, we demonstrated that sonogenetic-fPET could map the connectivity of this circuit, involving well-established brain regions such as the globus pallidus (GP), substantia nigra (SN), and Thalamus. fPET also recorded the activation of the contralateral limb muscle, demonstrating sonogenetics-fPET’s capability to identify the functional output of the neural circuit. By integrating acoustic hologram into the sonogenetic-fPET, we demonstrated its capability to flexibly probe different brain regions in the same animal. In summary, sonogenetic-fPET addresses the need for noninvasive and precise probing of the global neural network throughout the brain and body.

## Results

### Developing and Validating the Sonogenetic-fPET Technique

We developed the sonogenetic-fPET by integrating sonogenetics with whole-body fPET **(Fig. 1)**. This system involves a wearable ultrasound transducer for precise sonogenetic stimulation of specific brain regions, combined with a whole-body PET/CT scanner to capture resulting functional activity changes in the brain and body **(Fig. 1)**.

**Figure 1.**
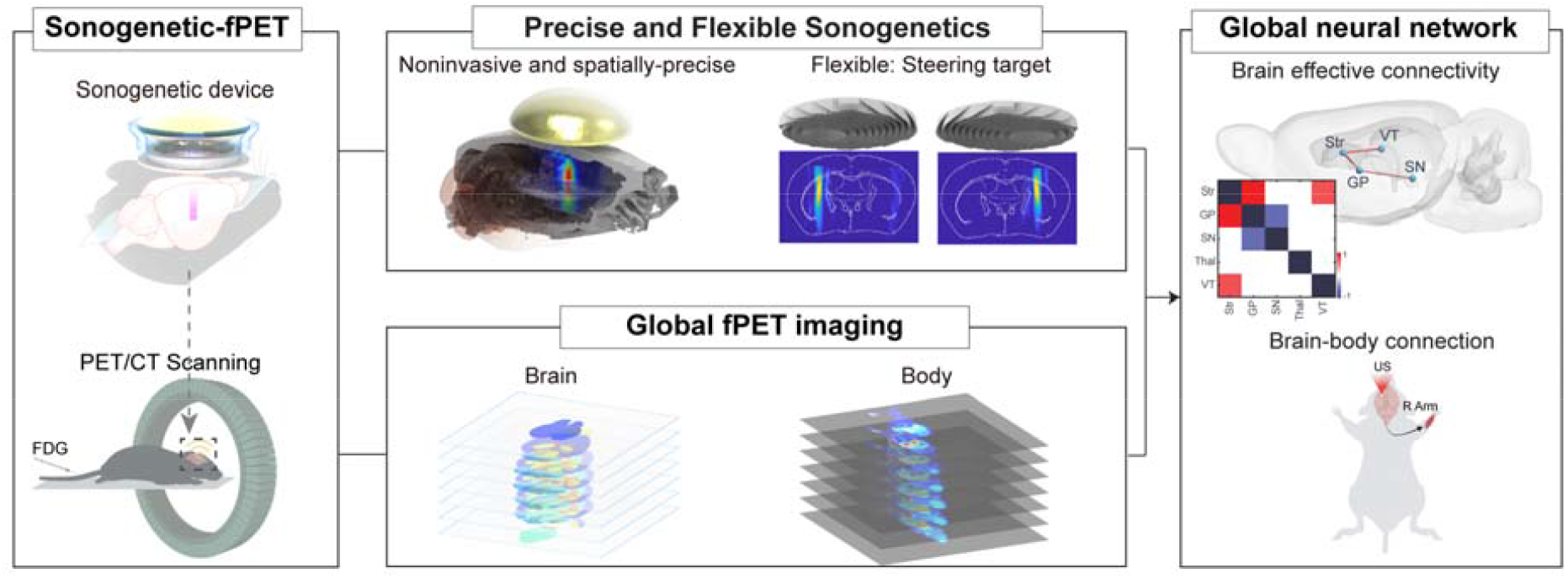
Overview of the sonogenetic functional positron emission tomography (sonogenetic-fPET) technique. The schematic depicts the hardware design, highlighting the features of sonogenetics and fPET. It also demonstrates the platform’s ability to map effective brain connectivity and brain-body interactions.

We developed a miniaturized wearable focused ultrasound transducer compatible with the PET scanner to deliver ultrasound stimulation to targeted brain regions. We validated its spatial focus using numerical simulations and experimental calibration in a water tank with an ex-vivo mouse skull (**Fig. 1**). The full width at half maximum (FWHM) of the ultrasound transducer was measured to be approximately 1.2□mm and 6□mm in the lateral and axial directions, respectively. We then placed mice with the wearable focused ultrasound transducer into the PET scanner and utilized [^18^F]-Fluoro-2-deoxy-2-D-glucose (FDG) as a radiotracer to map activation-induced metabolic increases in the brain and other organs, visualized as glucose uptake in the PET images. To identify brain activity changes in different regions, we registered the PET images to a standard brain atlas through a two-step process: registering PET images to anatomical CT images, followed by registration to the standard brain atlas using anatomical markers.

To validate the efficacy of the sonogenetic-fPET technique, we addressed two critical questions: (1) Can sonogenetics evoke spatially precise neuromodulation? (2) Can this evoked activity be detected by fPET? Sonogenetics was unilaterally targeted at the striatum, a core component of the basal ganglia circuit. Before sonogenetic stimulation, striatal neurons were engineered to express the ultrasound-sensitive TRPV1 ion channel via unilateral intracranial injection of AAV5-CaMKII-TRPV1 one month prior to stimulation. During the experiment, PET scanning started simultaneously with sonogenetic stimulation and FDG injection. Sonogenetic stimulation activated the targeted brain regions in TRPV1+ mice, demonstrated by increased FDG uptake at the target compared to its contralateral location (**Fig. 2A**). The dynamics of FDG activity from normalized fPET images at the targeted location showed enhanced activity (At 5 min: 0.08 ± 0.03 in sonogenetic group vs. -0.04 ± 0.02 in control; at 10 min: 0.15 ± 0.06 in sonogenetic group vs. -0.03 ± 0.06 in control) specifically during sonogenetic stimulation, with signals returning to baseline after sonication (**Fig. 2B**). No activation was observed in sham groups without ultrasound stimulation or TRPV1 expression. fPET results also demonstrated that sonogenetic-evoked brain activation is spatially precise, with FWHM of the activated region being approximately 1.2 mm along the anterior-posterior direction (**Fig. 2C**).

**Figure 2.**
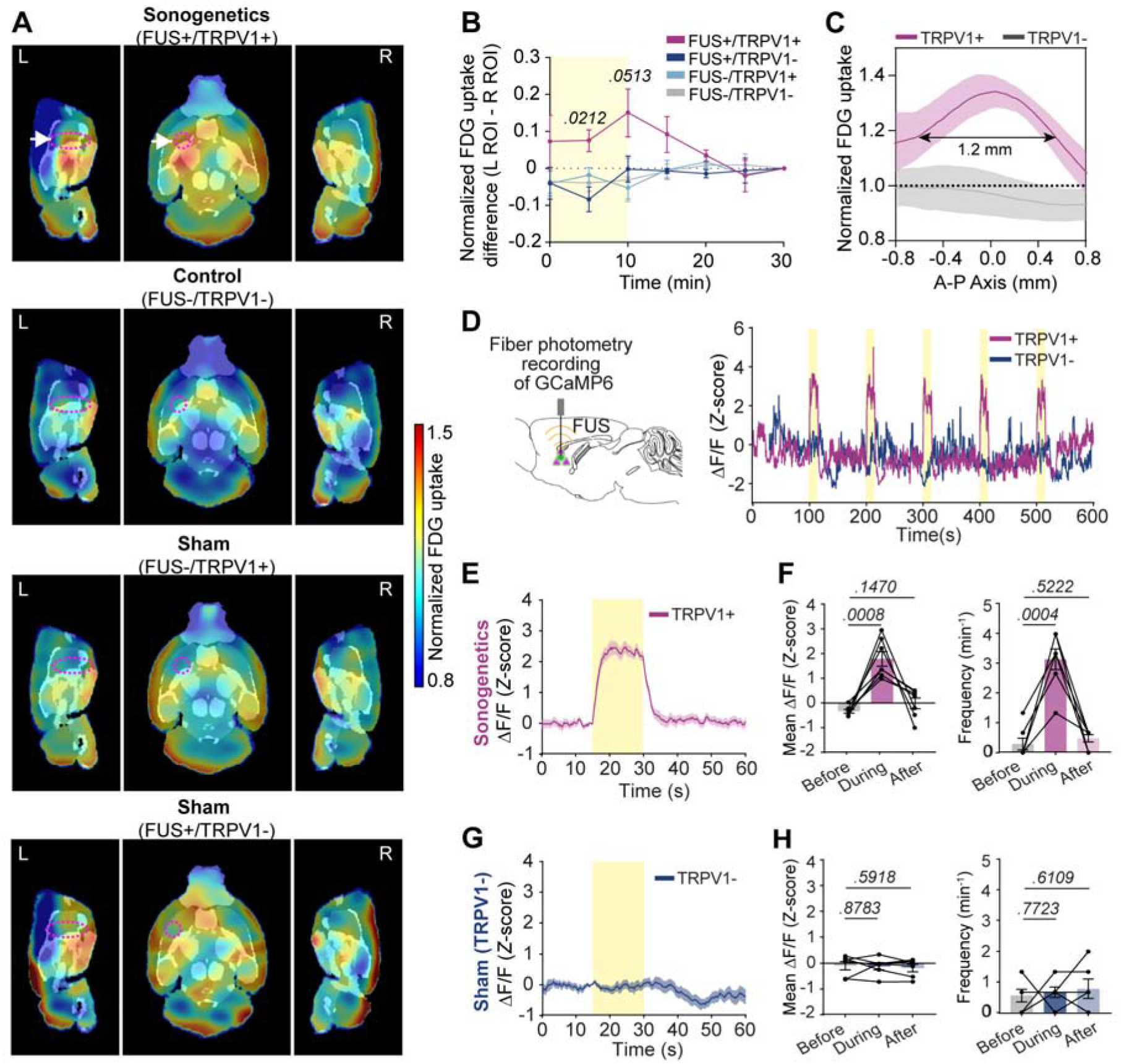
Sonogenetic-fPET is validated by measuring sonogenetically induced activation in the targeted brain region. **(A)** Averaged fPET images at the midpoint of focused ultrasound (FUS) stimulation (5 min), comparing different mouse groups: FUS+/TRPV1+ (sonogenetic group: FUS stimulation in TRPV1+ mice, n = 4), FUS-/TRPV1-(control group: no FUS stimulation in TRPV1-mice, n = 6), FUS-/TRPV1+ (sham group: no FUS stimulation in TRPV1+ mice, n = 9), and FUS+/TRPV1-(sham group: FUS stimulation in TRPV1-mice, n = 4). The color bar represents normalized FDG uptake. The images are from fPET during FUS stimulation, normalized to both the final scanning image after stimulation and the contralateral hemisphere, which did not receive FUS stimulation, to account for variations in FDG uptake across animals, which may arise from differences in body weight, blood glucose levels, and anaesthesia depth. Dotted circles indicate the FUS-targeted regions in the FUS+ groups and white arrows indicate the sonogenetic-evoked fPET signal. **(B)** Temporal quantification of the difference in normalized FDG uptake between the FUS-targeted region of interest (ROI) and the contralateral non-FUS site. **(C)** Spatial profile of normalized FDG uptake along the anterior-posterior axis at the FUS target site for the sonogenetic (TRPV1+) and control groups (TRPV1-). **(D)** Left: Schematic showing fiber photometry recording of neuronal Ca^2+^ activity in the striatum, targeted by FUS. Right: Representative Ca^2+^ signals for the sonogenetic (TRPV1+) and sham (TRPV1-) groups in response to FUS stimulation, with five FUS stimuli indicated by yellow bars. **(E)** Averaged Ca^2+^ signal for the sonogenetic group (TRPV1+, n = 7). **(F)** Mean ΔF/F and frequency of Ca^2+^ signals pre-, during, and post-FUS stimulation for the sonogenetic group. **(G)** Averaged Ca^2+^ signal for the sham group (TRPV1-, n = 6). **(H)** Mean ΔF/F and frequency of Ca^2+^ signals for the sham group (TRPV1-) pre-, during, and post-FUS stimulation. Statistical significance was determined using a paired two-tailed t-test for panels (F) and (H) and one-way analysis of variance for panel (B). p-values are indicated in the figures. Solid lines with shaded areas represent the mean ± s.e.m., and error bars indicate s.e.m.

To confirm that the evoked fPET signal was due to neuronal activation, we used fiber-photometry to record Ca^2+^ activity in sonogenetically stimulated neurons co-transfected with the Ca^2+^ reporter GCaMP6s and the sonogenetic actuator TRPV1. Sonogenetics evoked Ca^2+^ activity in targeted neurons, shown by a significantly elevated mean ΔF/F (from -0.33 ± 0.08 before stimulation to 1.80 ± 0.30 during stimulation) (**Fig. 2D–F**). In contrast, ultrasound stimulation alone in TRPV1-mice did not evoke significant neuronal activity (**Fig. 2G, H**). This observation aligns with the increased signal in fPET images in sonogenetically stimulated brain regions of TRPV1+ mice but not in TRPV1-mice, validating the capability of sonogenetic-fPET in spatially precise neuromodulation and recording.

### Sonogenetic-fPET Maps Effective Connectivity of the Basal Ganglia Circuit

We then examined the application of sonogenetic-fPET in mapping effective connectivity of the basal ganglia circuit. We activated the circuit via sonogenetic stimulation of the left striatum and derived the Z-score connectivity map from fPET images acquired during stimulation^33,35^. The map revealed activation in various brain regions associated with the striatum: the subthalamic nucleus (STN), ventral nucleus of the thalamus (VT), thalamus (Thal), the globus pallidus (GP) and the substantia nigra (SN) (**Fig. 3A**). These regions are well-documented as primary hubs within the basal ganglia circuit, receiving efferent projections from the striatum. Other brain regions not involved in the basal ganglia circuit, such as the hippocampus and olfactory bulb, showed minimal Z-score signal, suggesting the specificity of the sonogenetic-fPET. The Z-score connectivity map also aligned well with anatomical projection maps^36^, showing projections from the striatum to the GP, Thal, VT, and SN regions (**Fig. 3B**).

**Figure 3.**
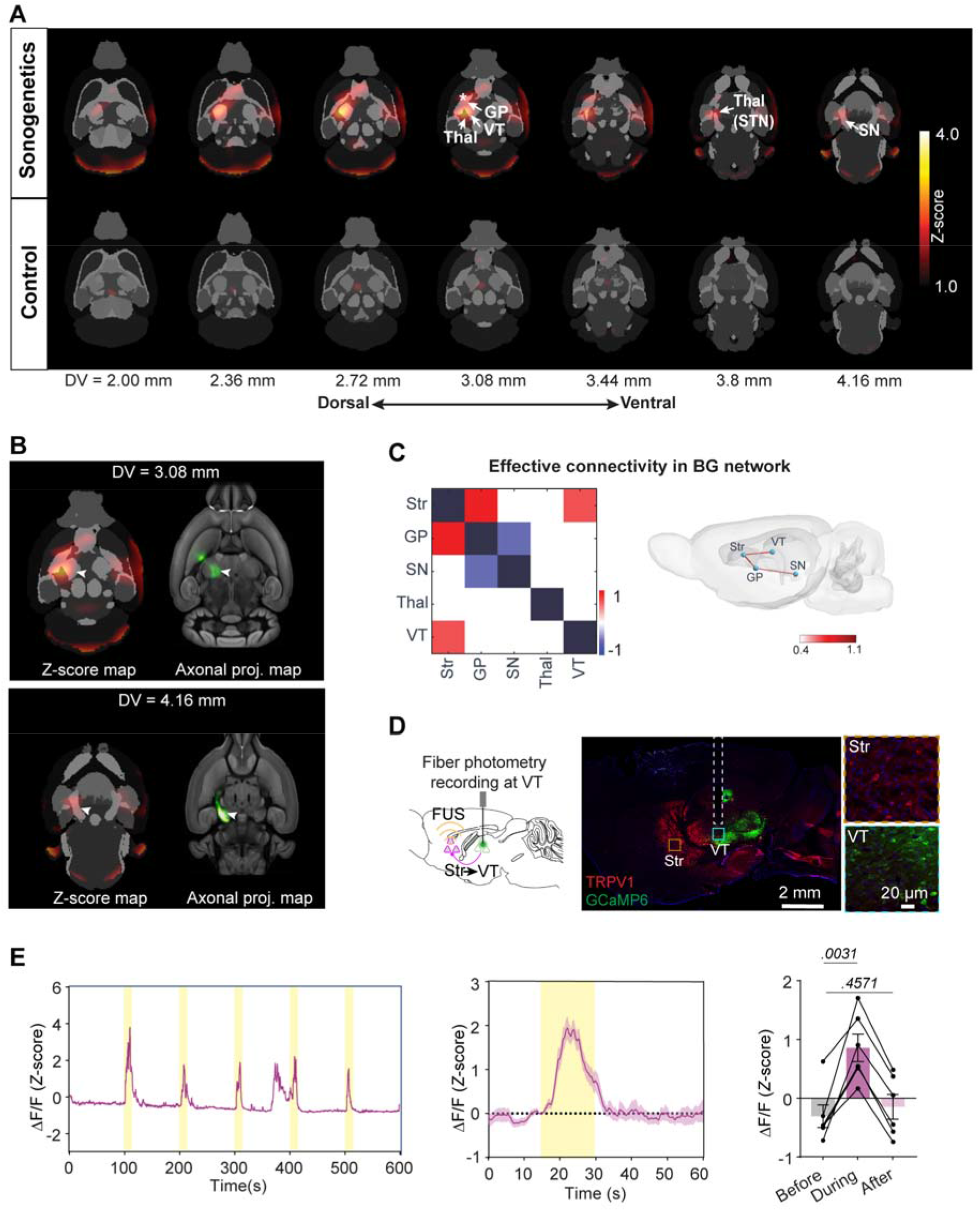
Sonogenetic-fPET maps the effective connectivity of the basal ganglia circuit. **(A)** Z-score connectivity maps corresponding to sonogenetic stimulation at the left striatum (top panel) and the control group (bottom panel). N = 4 mice for sonogenetic group and n = 6 for control group. **(B)** Comparison between the Z-score connectivity map and axonal projection patterns from the Allen Mouse Brain Connectivity Atlas at different dorsoventral (DV) depths (Experiment #113766038, https://connectivity.brain-map.org). **(C)** Effective connectivity map of the basal ganglia circuit and its corresponding network during sonogenetic stimulation at the striatum (Str) (p < 0.05). **(D)** Left: Schematic of the setup for simultaneous photometry recording of neuronal Ca^2+^ activity in the VT during sonogenetic stimulation at the striatum. Expression patterns of TRPV1 (red, immunofluorescence staining) and GCaMP6 (green) are shown. **(E)** Representative Ca^2+^ signals from the VT in response to sonogenetic stimulation at the striatum, along with the mean Ca^2+^ signal in the VT during sonogenetic stimulation. Sonogenetic stimulation periods are indicated by yellow bars. N = 6 mice. Solid lines with shaded regions represent mean ± s.e.m., and error bars indicate s.e.m. Statistical significance was assessed using a paired two-tailed t-test.

We then generated an effective connectivity network by integrating graph-theoretical analyses-based computational modeling with the sonogenetic-fPET results^37^. The connectivity analysis identified the striatum as the primary hub within the neural circuit, consistent with the experimental stimulation region (**Fig. 3C**). It revealed two connections: one projecting from the striatum to the GP and the SN and another projecting from the striatum to the VT (**Fig. 3C**). The former is consistent with the direct pathway of the basal ganglia circuit, which has been well studied^6,38^. To further validate the identified neural circuit connecting the striatum and VT, we recorded Ca^2+^ signals in the VT region via fiber photometry while sonogenetically stimulating striatal neurons. Sonogenetic activation of the striatum reliably provoked Ca^2+^ signals in the ipsilateral VT region, evidenced by an increased mean ΔF/F (from -0.31 ± 0.19 to 0.86 ± 0.23) (**Fig. 3D, E**). These results demonstrate that sonogenetic-fPET can map the effective connectivity of the basal ganglia circuit.

### Sonogenetic-fPET Probes Basal Ganglia Circuit Connection to Arm Muscles

Having demonstrated the capability of sonogenetic-fPET to map neural circuits, we investigated its ability to probe the connectivity of the neural circuits to peripheral organs, leveraging fPET’s unique capability to detect whole-body activity. Given the basal ganglia circuit’s role in motor control, we evaluated metabolic activity changes in peripheral limb muscles. Sonogenetic stimulation of the left striatum led to increased FDG uptake in the right forearm and palm muscles, contralateral to the targeted location (**Fig. 4A**). We observed higher FDG uptake in the contralateral forearm and palm muscles compared to the ipsilateral side (**Fig. 4B-D**; SUV = 0.63 ± 0.03 in the contralateral arm vs. 0.48 ± 0.04 in the ipsilateral arm at 5 min during stimulation). This uptake pattern was absent in sham groups without ultrasound stimulation or TRPV1 expression (**Fig. 4C, D**). These findings suggest that the function of the basal ganglia circuit in controlling motor behavior operates through its output to contralateral limb muscles. This observation aligns well with our previously reported behavior test results^16^, which showed a biased rotational behavior in one direction after sonogenetic stimulation of the striatum.

**Figure 4.**
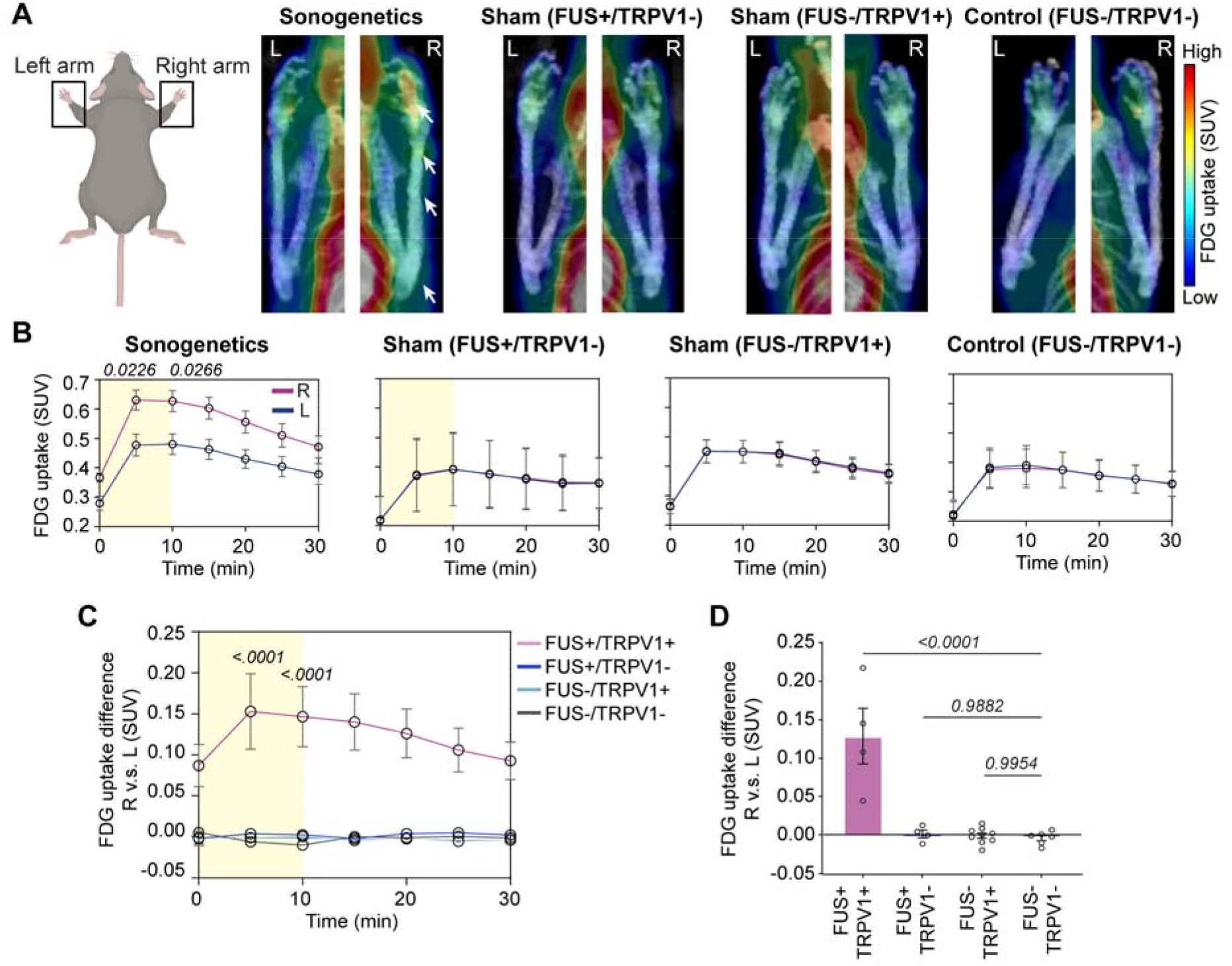
Sonogenetic-fPET probes the functional output of the basal ganglia circuit in contralateral limb muscles. **(A)** Representative fPET images showing FDG uptake activity in the left (L) and right (R) forearms during FUS stimulation across different groups, including FUS+/TRPV1+ (n = 4), FUS+/TRPV1-(n = 4), FUS-/TRPV1+ (n = 9), and FUS-/TRPV1-(n = 6). Arrows highlight areas of enhanced activity. **(B)** Temporal signal of FDG uptake for the right (R) and left (L) arms in different mouse groups. **(C)** Difference in FDG uptake between the right and left arms (calculated as FDG uptake in the right arm minus uptake in the left arm) in different mouse groups. **(D)** Average of FDG uptake difference between the right and left arms over the FUS stimulation period, with comparisons across groups. Yellow bars indicate the timing of sonogenetic stimulation. Error bars represent s.e.m. Statistical significance was determined using paired t-tests for panel (B) and one-way ANOVA for panels (C) and (D).

### Acoustic Hologram Enables Flexible Probing of Different Brain Regions in a Single Mouse

To improve the flexibility and throughput of sonogenetic-fPET, we manufactured a wearable focused ultrasound transducer using 3D-printed holographic lenses to flexibly target different brain regions within the same mouse by switching the holographic lenses^18,39^. We tested the performance of the holographic sonogenetics-fPET by alternately targeting the left and right striatum (**Fig. 5A**). Before the experiment, both the left and right striatum were engineered to express TRPV1 (**Fig. 5A**). fPET showed that the activation signal switched from the left striatum (1.13 ± 0.03 in the left vs. 0.9 ± 0.07 in the right) to the right striatum (1.21 ± 0.05 in the right vs. 0.93 ± 0.08 in the left) in response to ultrasound target steering (**Fig. 5B, C**), demonstrating that holographic sonogenetics can flexibly switch targeted brain locations within the same mouse. Moreover, fPET revealed that stimulation of the left striatum evoked higher activity in the right arm muscle (SUV = 0.62 ± 0.02) compared to the left arm (SUV = 0.52 ± 0.03). Shifting the ultrasound focus from the left to the right striatum evoked higher activity in the left arm than the right arm (SUV = 0.56 ± 0.05 in the left arm vs. SUV = 0.49 ± 0.04 in the right arm) (**Fig. 5D**). These findings demonstrated the capability of hologram-mediated sonogenetic-fPET to flexibly probing different brain regions in a single mouse.

**Figure 5.**
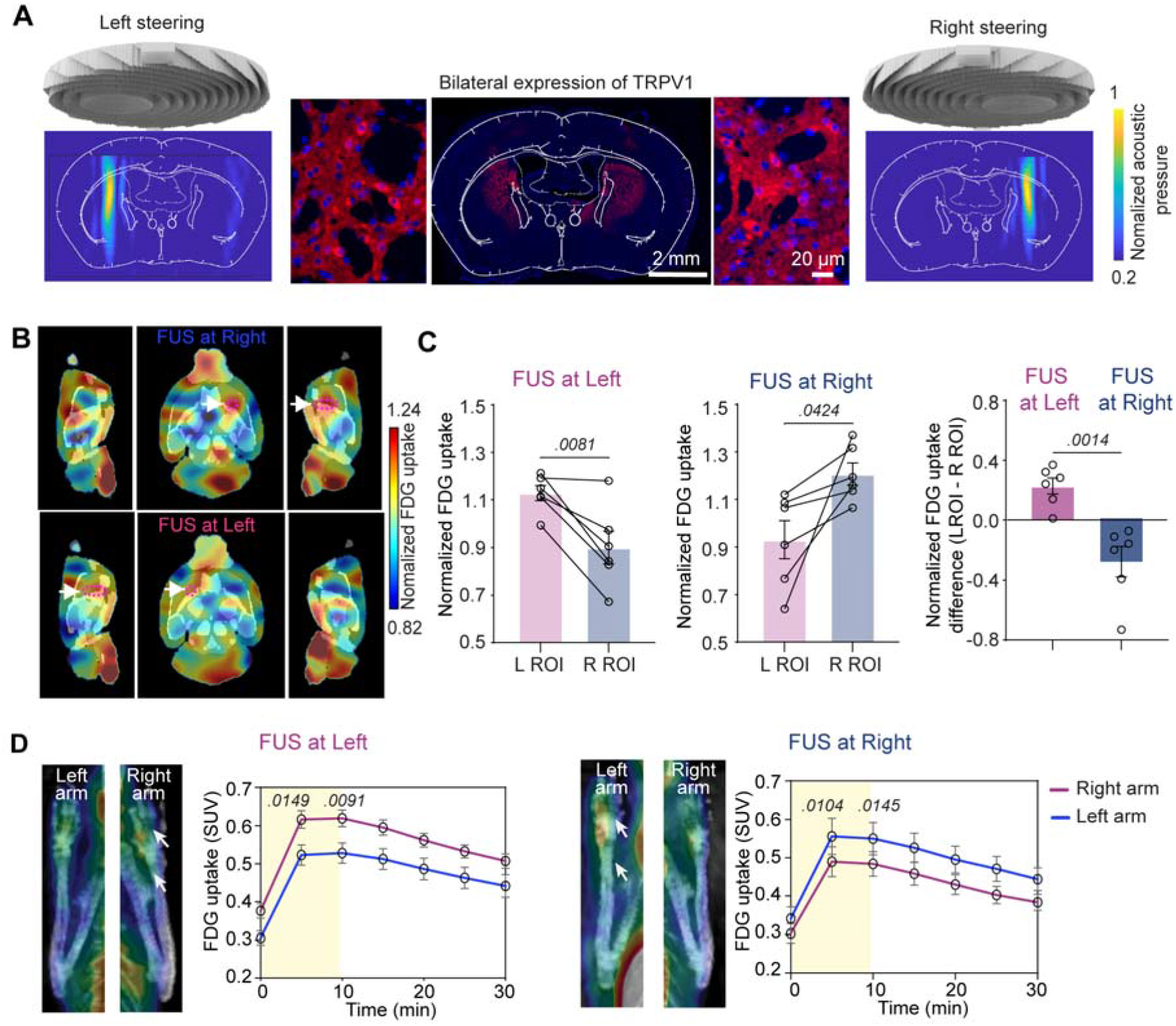
Acoustic hologram enhances the flexibility of sonogenetic-fPET for targeting different brain regions within the same animal. (**A**) Acoustic patterns produced by the acoustic hologram, designed to target either the left or right striatum—both expressing TRPV1. **(B)** Averaged fPET images across all mice undergoing holographic sonogenetic stimulation, targeting either the left (upper panel) or right (lower panel) striatum. Color bars represent the normalized FDG uptake. Dotted circles and arrows highlight the FUS-targeted regions. N = 6 mice. **(C)** Spatially averaged signal of the normalized FDG uptake in the left and right ROIs (FUS targeted region and its corresponding non-targeted contralateral site) for mice receiving holographic sonogenetic stimulation on either the left (FUS at Left) or right (FUS at Right) striatal regions. The right panel shows the difference in normalized FDG uptake between the left and right striatal ROIs during holographic sonogenetic stimulation at the corresponding side. **(D)** Representative fPET images of the left and right forearms during holographic sonogenetic stimulation targeting either the left or right striatum, along with corresponding temporal signal of FDG uptake in the right and left forearms. Error bars represent s.e.m. Statistical significance was determined using a paired two-tailed t-test.

## Discussion

In this study, we developed sonogenetic-fPET by integrating sonogenetics with FDG-fPET to map global networks throughout the brain and body evoked by modulation of genetically and spatially targeted neural populations. Using the basal ganglia circuit as a model, we demonstrated that this technique enables the mapping of brain neural circuits and the activation of downstream effector tissues. The incorporation of an acoustic hologram further enhances the flexibility of sonogenetic-fPET, enabling flexible switching of the targeted brain regions within a single mouse.

We validated the effectiveness of sonogenetic-fPET by mapping the connectivity of the basal ganglia circuit. Previous studies have shown that the direct pathway in the basal ganglia facilitates movement through projections from the striatum to the GPi and SN, leading to disinhibition of the thalamus^6,40,41^. In contrast, the indirect pathway inhibits movement by projecting from the striatum to the GPe, then to the subthalamic nucleus (STN), and ultimately to the GPi and SN, resulting in thalamic inhibition^42^. Our findings align with these models, as we identified functional connections from the striatum to the GP and SN, suggesting activation of the direct pathway via sonogenetic stimulation. While we did not specifically target the direct pathway by using D1-Cre mice as reported in previous studies ^6^, previous studies found that sonogenetics with the CaMKII promoter biases activation towards this pathway, leading to contralateral rotational behavior^16^ —consistent with our observations. Although we did not detect significant STN activation in our connectivity network, likely due to its small size and the limited spatial resolution of fPET, a Z-score of 2.49 in the connectivity map suggests some level of activity change in response to sonogenetic stimulation. This limitation is inherent to PET imaging, which has a limited spatial resolution. We also observed elevated glucose uptake in the contralateral limb muscles, indicating a functional connection between the basal ganglia circuit and the limb muscles, consistent with existing literature^6,40,41^. Our data validate the effectiveness of sonogenetic-fPET in mapping basal ganglia circuit connectivity and its downstream effects on limb muscles.

Integration of the acoustic hologram into the sonogenetic-fPET technique unlocks new capabilities for mapping various neural circuits within the same mouse. The acoustic hologram allows precise and flexible targeting of the ultrasound beam to different brain regions, enabling the study of multiple circuits in the same animal. We demonstrated this by switching the target between the left and right striatum using different hologram lenses, successfully evoking distinct brain region activation as designed. This capability, unique to sonogenetics, represents a significant advancement over conventional implant-based neuromodulation technologies like optogenetics and electrical stimulation. The holographic ultrasound transducer also has the potential to generate arbitrarily 3D acoustic patterns^18,39,43^. These patterns could further enhance sonogenetics’ ability to target multiple brain regions simultaneously or “paint” activation patterns within the brain, potentially advancing our understanding of region-to-region interactions and refining sub-brain region connections. By combining its flexible targeting with global mapping capabilities, sonogenetic-fPET has the potential to greatly accelerate neural circuit discovery and enhance our understanding of brain function.

Sonogenetic-fPET expands the toolbox for neuroscience research by combining precise “writing” (sonogenetics) and global “reading” (fPET). Sonogenetics enables targeted activation of spatially and genetically defined neurons in the whole brain, while fPET captures metabolic activity in both the brain and body. This technique addresses the limitations of opto-fMRI and provides a complementary tool to existing techniques^1–3^. Optogenetics typically requires invasive implants, while sonogenetics can target deep brain regions without any implants. fMRI can map neural circuits within the whole brain but provides limited information about their connections to peripheral organs and tissues. This gap limits our understanding of the brain-body connection, which is crucial for a comprehensive understanding of brain function. Emerging technologies like multifunctional microelectronic fibers can modulate and probe the brain-gut axis but require invasive implants, limiting their application^44^. Other approaches, such as whole-mouse clearing and imaging technologies, provide detailed anatomical maps but lack functional connectivity information and require sacrificing the animal^45^, limiting their utility in longitudinal studies. Sonogenetic-fPET overcomes these limitations by noninvasively probing intact neural networks. In addition, focused ultrasound techniques and PET imaging have already been used in humans, laying the groundwork for future applications of sonogenetic-fPET in humans.

In conclusion, sonogenetic-fPET addresses the need for noninvasive, precise neural circuit probing, and global neural network mapping. This noninvasive, implant-free approach allows for studying intact neural networks throughout the brain and body with minimal disruption to the intrinsic natural circuitry. The integration of acoustic hologram technology further enhances its flexibility, allowing for targeted probing of different neural circuits within the same animal. This technique has the potential to accelerate neural circuit discovery and enable the development of circuit-specific precise treatments for brain diseases.

## Methods

### Animals

Female C57BL/6NCrl mice (Charles River Laboratories) aged between 6-9 weeks were randomly assigned to different groups for this study. The mice were housed at the animal facility of Washington University School of Medicine, where they were kept in a controlled environment with a temperature of 23-26°C and humidity levels between 35-65%. The mice were also provided with a standard chow diet and subjected to a 12-hour light/dark cycle. All animal studies conducted in this research were reviewed and approved by the Institutional Animal Care and Use Committee (IACUC) of Washington University in St. Louis. These studies were carried out in accordance with the National Institutes of Health Guidelines for Animal Research, and the assigned animal protocol number was 21-0187.

### Virus injection

In this study, we overexpressed TRPV1 in striatal neurons by injecting AAV9-CaMKII-TRPV1-p2A-mCherry (0.2 µl, 4.4 × 10^13^ vg/ml) into the left dorsal striatum (AP = ∼0.0 mm, ML = ∼2.0 mm, DV = ∼3.0 mm), following a previously described intracranial injection procedure^19^. To establish a sham control group, AAV9-CaMKII-mCherry (0.6 µl, 1.8 × 10^13^ vg/ml, Addgene, #114469-AAV9) was injected to ensure a similar viral vector number. Before the viral injection, mice were anesthetized with 1.5% isoflurane and administered subcutaneous buprenorphine (Buprenex, 0.1 μg/g body weight) for analgesia. The AAVs were delivered into the striatum at a rate of 0.69 µl/min using a microinjector (Nanoject II; Drummond Scientific) after creating a small (∼0.7 mm) cranial burr hole. The injection needle was withdrawn slowly at a rate of approximately 3.0 mm/min one-minute post-injection, and the scalp was sutured. Mice were then allowed to recover on a heating pad. To record neuronal calcium activity during ultrasound sonication, AAV5-Syn-GCaMP6s (0.3 µl, 1.0 × 10^13^ vg/ml, Addgene, #107790-AAV9) was injected into the same brain region using the same procedures. Additionally, to test the capability of the hologram transducer to steer targets between the left and right striatum, the virus was injected bilaterally into the dorsal striatum.

### FUS stimulation

We used miniaturized wearable ultrasound transducers for delivering ultrasound stimulation, as previously reported ^16^. These transducers were constructed using a concave-shape lead zirconate titanate (PZT) ceramic resonators (DL-43, DeL Piezo Specialties, FL) with a center frequency of 1.5 MHz, an aperture of 10 mm, and a geometric focal depth of 10 mm. Additionally, we designed and applied an ultrasound holographic transducer using a previously described method to steer the ultrasound target to either the left or right striatum ^39^. Briefly, the hologram transducer consisted of a plane PZT material with a 13 mm aperture and a 3.2 MHz center frequency, along with a 3D-printed hologram lens designed by time-reversal methods ^39^, without the binary process in this study. The transducers were calibrated using a hydrophone (HGL-200, Onda) in degassed and deionized water, both with and without an ex-vivo mouse skull. FWHM of the concave transducer was 1.2 mm and 6 mm in the lateral and axial directions, respectively. And FWHM of the holographic transducer was 0.5 mm and 5.5 mm.

To attach the wearable ultrasound transducer to the mouse head, we used a plug-in design onto a baseplate that was glued onto the mouse skull approximately 3 weeks after virus injection. Specifically, mice were anesthetized and head-fixed using a stereotactic frame, and the baseplate was glued onto the top of the mouse skull using adhesive cement, with the center of the baseplate aligned to the center of the virus injection site (AP = ∼0.0 mm, ML = ∼2.0 mm). Animals were given at least 3 days to recover and adapt to the baseplate post-surgery. Prior to each ultrasound stimulation experiment, the wearable ultrasound transducer was filled with degassed ultrasound gel (Aquasonics) and plugged into the baseplate on the mouse head. The ultrasound stimulation parameters were set to acoustic pressure = 1.3 MPa, a frequency of 1.5 MHz (concave shape transducer) or 3.2 MHz (holographic transducer), 40 ms pulse duration (duty cycle = 40%), 10 Hz pulse repetition frequency, 15 s stimulus duration, 100 s inter-stimulus interval, and a total of 6 stimuli.

### PET/CT scanning

PET scanning was conducted to image the brain and body activity of the mice in response to ultrasound stimulation. Animals were first anesthetized via intramuscular injection of a ketamine (0.10 mg/g body weight) and xylazine (0.01 mg/g body weight) mixture. The transducer was then plugged onto the base plate (taking ∼15mins). The anesthetized mouse, with a wearable ultrasound transducer attached, was then placed into the chamber of the Inveon CT/PET scanner (Siemens, Malvern, PA), where a CT scan was performed to determine anatomical placement. Next, FDG (∼250-300 µCi, 100 µL volume) was injected into the tail vein catheter of the mouse using a 0.300 cc, 29 Gauge x ½” insulin syringe, while ultrasound stimulation was simultaneously initiated with PET scanning. The injected dose of FDG was precisely measured by counting it in a Capintec dose calibrator before and after the injection. PET acquisition was captured for 30 minutes, and the acquired PET images were corrected for attenuation, scatter, normalization, and camera dead time, then co-registered with CT images. During the entire procedure, the mice were kept warm using a lamp. The voxel size of the PET and CT images was 0.8 mm × 0.8 mm × 0.8 mm and 0.2 mm × 0.2 mm × 0.2 mm, respectively.

### fPET image analysis

The FDG uptake in the fPET images was calculated to the standardized uptake value (SUV) by the algorithm 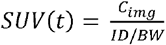, where *c*_*img*_ refers to the image-derived radioactivity concentration at time *t* post-injection; ID and BW refer to the total injected dose and body weight, respectively. To locate the FUS target and different brain regions in fPET images, anatomical CT scans and fPET images were registered to the 3D brain atlas in the following steps. First, CT images were registered to fPET images using ITK-SNAP (version 4) software with only the translation process, following the resampling of the CT voxel size to match the PET voxel size (0.8 mm × 0.8 mm × 0.8 mm). Next, CT images were registered to the standard brain atlas ^36^ by matching the key anatomical skull markers in the CT images with the brain atlas, using scaling, translation, and rotation processes in the same software. The transformation matrix was then exported to apply to the fPET image for its registration to the brain atlas. To visualize the FUS-induced signal change, the last fPET frame acquired at 30 minutes (20 minutes after FUS stimulation) was subtracted from the other image stacks to generate normalized fPET images. To visualize the group effect, all the normalized fPET images from the same group were averaged together after they were registered to the standard brain atlas.

To examine the dynamics of brain activity induced by sonogenetics, we calculated the spatial average of FDG uptake values in both FUS-targeted ROI and the contralateral non-targeted ROI over time in the normalized fPET images. FUS-targeted ROIs were selected based on the estimated virus injection site, which is also the FUS-targeted location with the size estimated by the calibrated FWHM of the FUS beam. The difference in average FDG uptake between the FUS-targeted ROI and the corresponding contralateral ROI of the same size represented the FUS-induced brain activity change. Additionally, we quantified the FDG uptake in the forearm muscle within a cylindrical shape (∼4 mm diameter) manually selected to cover the palm and the forearm muscle identified in the CT images.

We also assessed the spatial resolution of sonogenetics by calculating the FWHM of the normalized fPET signal in the FUS-targeted brain region. Specifically, we derived a differentiated FDG uptake image by subtracting the baseline activity (averaged signal from the contralateral hemisphere without FUS stimulation) from the normalized PET images and then calculated the FWHM centered at the maximum pixel at the FUS-targeted site on the horizontal plane, which is approximately perpendicular to the FUS incident beam.

### Connectivity map

The Z-score connectivity map was generated from the normalized fPET images during FUS stimulation period (at 5 min). To compute Z-scores, the normalized PET images were first adjusted by dividing by the mean of the contralateral side to account for differences in mouse metabolism and baseline activity. Then, the mean and standard deviation of all voxels across all scans in the treatment group were calculated. The standardized Z-score was computed on a voxel-by-voxel basis by subtracting the mean from the normalized activity level and dividing by the standard deviation^33,35^. This process was repeated for all scans in each FUS treatment group. The effective connectivity map and network were generated using the open-source software GraphVar, following the published protocol^37^.

### Fiber-optic photometry recording

We investigated ultrasound-evoked neuronal activities in the ultrasound-targeted brain region and downstream VT region by recording GCaMP signals using fiber photometry. Three weeks after virus injection, a fiberoptic cannula (MFC_200/245-0.37_6mm_ZF1.25_FLT, Doric Lenses Inc.) was implanted approximately 200 µm above the injection site through the hole of the baseplate used for virus injection. This design allowed for the alignment of the ultrasound focal region and the tip of the cannula within the targeted striatum. To record GCaMP signals at the VT region, a baseplate with a hole at 2 mm off-center was implanted, and the cannula was inserted through this hole into the brain. This plate design allowed for ultrasound stimulation through the center of the plate aligned with the striatum while photometry recording was at the VT region. The cannula and baseplate were fixed and stabilized with adhesive cement, and the mice were allowed to recover for one week.

Wearable ultrasound transducers were built for simultaneous ultrasound stimulation and fiber photometry recording by drilling a 2 mm diameter hole for inserting the optical fiber ^18,19^. The hole was located at the center of the transducer for recording FUS-targeted striatum neurons and 2 mm posterior to the target site for recording signals in the downstream VT region. The transducer was plugged into the baseplate, and a mating sleeve was passed through the hole of the transducer to connect the stainless-steel ferrule to the implanted cannula. The fiber transmitted excitatory blue light (wavelength: 470□nm, power: 4%) and collected GCaMP6-emitted photons. GCaMP6 signals were collected, digitized, and measured with a photometry system (FP3002, Neurophotometrics) in synchronization with ultrasound stimulation. Photometry data were acquired before, during, and after stimulation, and an open-source software, Bonsai (v2.7), was used to acquire the photometry data and the synchronization signal from an LED. The acquired data were processed using a method adapted from a previously reported method^19^. Data were first de-bleached using a high pass filter and converted to a Z-score using the MATLAB “zscore” function, and the mean of the Z-score was calculated before FUS (the 15 s period right before FUS stimulation), during FUS (15 s during FUS stimulation), and after FUS (the 15 s period right after FUS stimulation ended).

## Statistical analysis

Statistical tests were conducted using GraphPad (Prism). Data were analyzed using one-way ANOVA and student’s t-test. Statistical differences were considered significant whenever *P* < 0.05. *P*-values are indicated in the figures. All the graphs presented the results as standard error of the mean.

## Reporting Summary

Further information on research design is available in the Nature Research Reporting Summary linked to this article.

## Data availability

We have uploaded the design file for the holographic transducer to GitHub: https://github.com/ChenUltrasoundLabWUSTL. Additionally, the plasmid is available through Addgene (Plasmid #200829). All other data are included in the manuscript. Source data for the figures will be provided in this paper. The mouse brain atlas used in this study is available in the Allen Brain Atlas.

## Author contributions

Conceptualization: HC, YYang. Experiment design: HC, YYang. Experiment implementation: YYang, CW, YYue, ZH, HC. Result Investigation: YYang, HC, CW, IW. Visualization: YYang, HC, CW, IW, LX. Funding acquisition: HC. Project administration: HC. Supervision: HC, YYang. Writing – original draft: HC, YYang, CW. Writing – review & editing: HC, YYang, CW, YYue, HC.

